# O mouse, where art thou? The Mouse Position Surveillance System (MoPSS) - an RFID based tracking system

**DOI:** 10.1101/2020.11.13.379719

**Authors:** Anne Habedank, Birk Urmersbach, Pia Kahnau, Lars Lewejohann

## Abstract

Existing methods for analysis of home cage based preference tests are either time consuming, not suitable for group management, expensive and/or based on proprietary equipment that is not freely available. For this reason, we developed an automated system for group housed mice based on radio frequency identification: the Mouse Position Surveillance System (MoPSS). The system uses an Arduino microcontroller with compatible components, it is affordable and easy to rebuild for every laboratory. The MoPSS was validated using female C57BL/6J mice and manual video comparison. It proved to be accurate even for fast moving mice (up to 100 % accuracy after logical reconstruction), and is already implemented in several studies in our laboratory. Here, we provide the complete construction description as well as the validation data and the results of an example experiment. This tracking system will allow group-based preference testing with individually identified mice to be carried out in a convenient manner, creating the foundation for better housing conditions from the animals’ perspective.

## 1 Introduction

Preference tests are increasingly used to improve the housing and living conditions of laboratory animals. Such test procedures allow the animals’ point of view to be directly involved in the refinement process. In order to get a meaningful impression of the choices made, the tests should largely reflect normal laboratory conditions and allow to record the choice behaviour without interference by an experimenter. This is at best realized using home-cage-based preference tests (Habedank et al., 2018). For mice, the apparatus for such a choice test usually consists of two (Kawakami et al., 2012; Kirchner et al., 2012; Loo et al., 2005) or more (Ago et al., 2002; Godbey et al., 2011; de Weerd et al., 1997) connected cages, directly connected via tubes or with a center cage. Animals are given continuous access to the options presented in each cage. In order to measure preference, either the nest position (Loo et al., 2005; Baumans et al., 2002) or the time the animals spent in the compartment (Godbey et al., 2011; Kawakami et al., 2012; Blom et al., 1992;Kirchner et al., 2012; Freymann et al., 2015; Freymann et al., 2017) is then monitored and regarded as the favored one (Habedank et al., 2018).

Thus, home cage based preference tests are based on binary or multiple choices, and they are designed to rank the preferences, not to assess the strength of preference or the “demand” for this resource (Kirkden and Pajor, 2006). In this manner the preference of mice was already investigated regarding bedding material (Kirchner et al., 2012; Blom et al., 1996), the provided amount of it (Freymann et al., 2015; Freymann et al., 2017), nesting material (Ago et al., 2002; de Weerd et al., 1997), shelters (Loo et al., 2005), cage-change interval (Godbey et al., 2011), ventilation (Baumans et al., 2002; Krohn and Hansen, 2010), temperature (Gaskill et al., 2009; Gaskill et al., 2011; Gaskill et al., 2012) and environment colour (Kawakami et al., 2012). Further husbandry conditions which to our knowledge are not yet fully investigated in this manner are, e.g., brightness, humidity and different items of enrichment such as structural elements or equipment for active engagement.

When conducting a home cage based preference test, it can be distinguished between the active (dark) and the inactive (light) phase to analyse the data (Lewejohann and Sachser, 2000; Freymann et al., 2015). This is especially important if the tested cage conditions are predominantly associated with active (e.g., running wheel) or inactive behaviour (e.g., nesting material). Social species of laboratory animals such as mice are usually kept in groups. Social conditions are likely to influence the choice of individual mice as for example, the sleeping temperature might be influenced by the presence of other animals. Thus, generally speaking, animals that are living in groups under normal laboratory conditions should also be tested in groups. However, measuring the preference of a group of mice is a far greater challenge than measuring singly housed mice, and thus, many of the preference studies investigated individual mice instead of groups (Blom et al., 1992; Blom et al., 1996; Kawakami et al., 2007; Kawakami et al., 2012; de Weerd et al., 1997). When testing groups (Kirchner et al., 2012; Freymann et al., 2015; Freymann et al., 2017; Godbey et al., 2011; Gaskill et al., 2011; Gaskill et al., 2009; Gaskill et al., 2012), individuals in one group can influence each other (Loo et al., 2001; Valsecchi and Galef, 1989; Shemesh et al., 2013), so that the results from one group might have to be counted as a single unit. More recent advances in statistical methods allow including “group” as a random factor in the model, but still the total number of animals might have to be increased to account for such group effects.

Of the available methods to analyse a home cage based preference test, most do not carry the capability to sufficiently cope with implicit challenges of choice tests. For example, monitoring only the nest position (Baumans et al., 2002; Loo et al., 2005) causes little costs with regard to equipment and time, but provides mainly information on where the mice spent their inactive time and thus does not reflect temporal distribution of individual preferences. The most common analysis of home cage based preference tests is therefore done by video recordings (Godbey et al., 2011; Kawakami et al., 2007; Ago et al., 2002; Gaskill et al., 2009; Gaskill et al., 2011). However, video analysis is very time consuming, especially when it is necessary to distinguish between individuals. For this reason, some research groups only analyse part of the recordings instead of a continuous tracking (every 5 min: Kawakami et al., 2007; every 10 min: Gaskill et al., 2009; Gaskill et al., 2011; Gaskill et al., 2012; every 60 min: Godbey et al., 2011), whereby the time saving is at the expense of the accuracy of the measurement. Analysis of the videos in a more automated manner by using video tracking software (Nath et al., 2019; Rao et al., 2019; Noldus et al., 2001) is by now not advanced enough to ensure decent tracking of individual mice in the test environment. However, there are other techniques which allow automated tracking: For example, in the connecting tunnels light barriers can be implemented to record whenever an animal changes cages (Blom et al., 1992; Blom et al., 1996). This method allows easy continuous tracking without much analysis effort. However, this approach is not suitable for group housing because aside from lacking individual detection, the determination of direction of passages is erroneous if sensors can be triggered by more than one animal. Similar problems would also arise if using digital scales below the cages combined with an automated tracking program (Krohn and Hansen, 2010).

To combine automated and individual detection, telemetry can be used by either implanting a rather large, battery-powered transponder (Kawakami et al., 2012) or injecting a smaller, passive transponder for radio-frequency identification (RFID) (Kirchner et al., 2012; Freymann et al., 2015; Freymann et al., 2017). The latter method is also very commonly used not just for choice tests but to record general activity patterns of mice (Freund et al., 2013; Bains et al., 2016; Weissbrod et al., 2013; de Chaumont et al., 2019), rats (Redfern et al., 2017) and birds (Bridge et al., 2019). But although there have already been automated tracking systems described, those are either expensive (Linnenbrink and von Merten, 2017; Bains et al., 2016), use non-implantable transponders (Bridge et al., 2019), can only detect animal species moving slower than mice (birds in nest boxes: Bridge et al., 2019) or they are based on proprietary equipment that is not freely available (Tsai et al., 2012). Thus, for a home cage based preference test with group housed mice, a reliable, time and cost efficient analysis method is still missing. (An overview of the described methods so far and their advantages and disadvantages is summarized in Table 1.) For this reason, we developed an automated system based on RFID which is affordable, easy to (re)build and suitable for individual tracking in group housed mice: the Mouse Position Surveillance System (MoPSS). It consists of an Arduino MKR WIFI 1010 microcontroller and two RFID controllers with two antennae. In order to read an RFID signal the transponder has to stay within the electromagnetic field of the antenna for around 30 ms. Mice are capable of very fast movements and can reach up to 18.0 m/min without training on a treadmill (Billat et al., 2005), 23-31.8 m/min after training (Hollinski et al., 2018), 67 m/min on a running wheel (Bono et al., 2006) and possibly even higher velocities during short sprints and jumping. Therefore, additional barriers were added in the connecting tube between the cages in order to slow down the movements in the vicinity of the antennae. Here, we provide the experimental validation of the system with a group of 7 weeks old female C57BL/6J mice as well as the complete implementation description: To facilitate the rebuilding of the MoPSS in other laboratories, we supply the construction plan, the Arduino code and the 3D print design of the barriers. We also describe an additional analysis method for the data which uses logical reconstruction to further improve the obtained data. With the help of this paper, the MoPSS can be rebuilt by any laboratory and/or altered with regard to alternative research questions (for example other species).

**Tab. 1:**
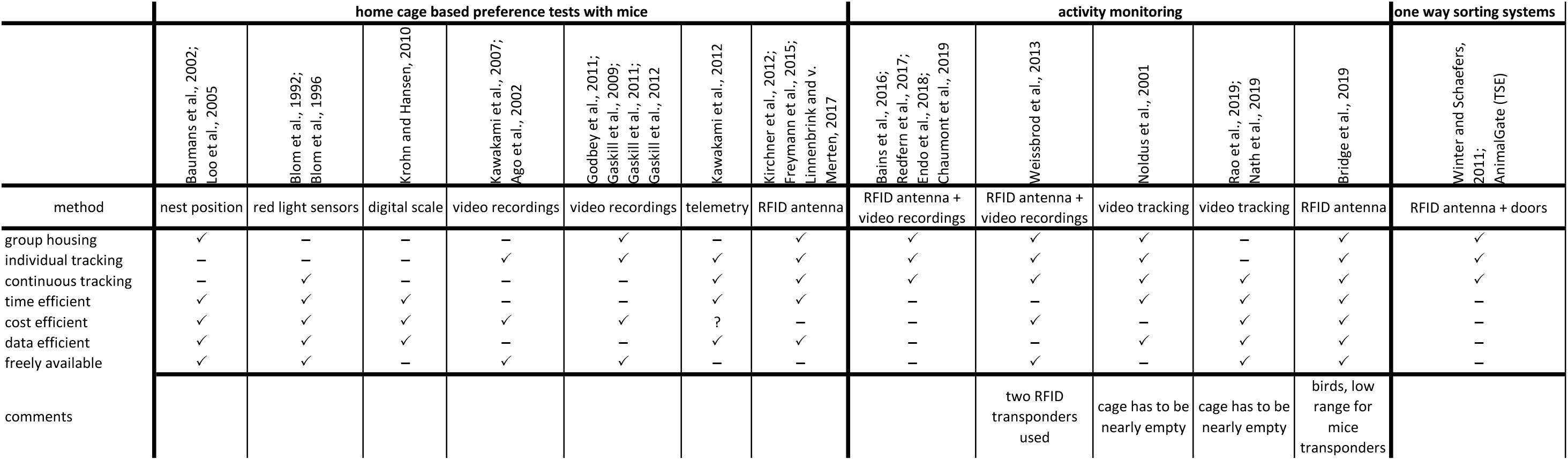
Described methods usable for home cage based preference tests. Methods are sorted for their purpose (used in home cage based preference test, used for activity monitoring but in general applicable for preference tests and used as a one way sorting mechanism, which would either have to be reprogrammed, or of which two would have to be used, for each direction one). The evaluation of the systems is done based on the named papers and what the authors described there. We did not add speculations, what might be additionally possible with these systems (e.g., whether group housing or tracking of individuals would have been feasible).

## 2 The Mouse Position Surveillance System (MoPSS)

### 2.1 General Principle

The basic experimental setup consists of two cages which are connected by a perspex tube (40 mm in diameter) passing two RFID antennas (see Fig. 1). As the system relies on RFID, all animals need to have an RFID transponder implanted. We recommend placing it under the skin in the neck region. For best reading performance the transponder must be implanted lengthwise (rostrocaudal). When a mouse moves through the tube and enters the magnetic field emitted by the RFID antenna, the transponder is read and the transponder number, antenna number, and current timestamp is saved onto a microSD card (32 GB). For the analysis, a mouse detected at the left RFID antenna is counted as being in the left cage, and a mouse detected by the right RFID antenna is counted as being in the right cage. It is possible to subtract the transition duration so as to not add it to one of the cages, however, as mice usually pass very quickly through the tube, we argue that the time passage time is neglectable. The main challenge while developing the apparatus was that mice were too fast for the RFID detectors, i.e., spending less time than necessary within the read range during the read cycle. In addition, if multiple mice were in the range of the same antenna interference led to poorer detection as well. Therefore, we added two barriers inside the connecting tube each obstructing approximately 40% of the tubes’ diameter and thereby forcing the mice to slow down in the vicinity of the antennae while passing the barriers.

**Fig. 1:**
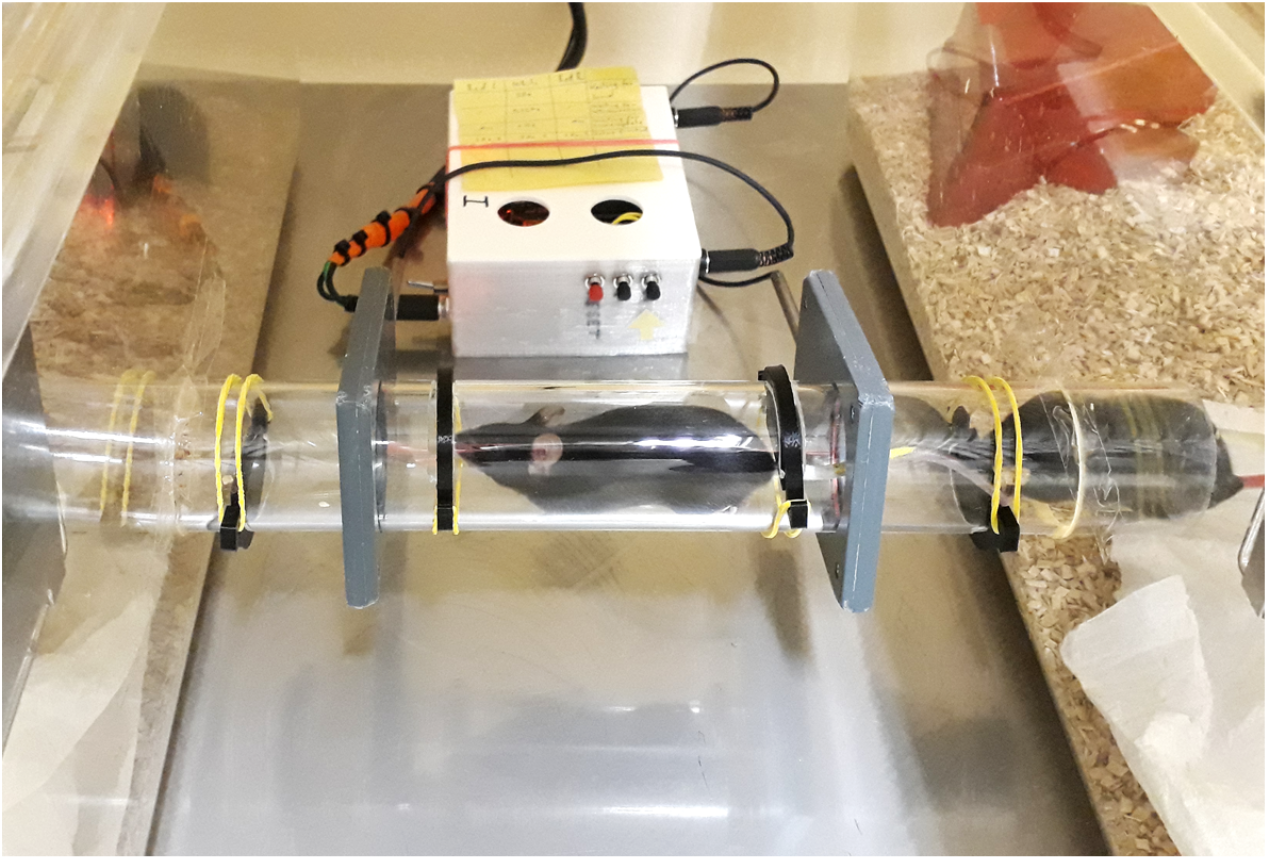
Setup of a home cage based preference test using the MoPSS. Two cages are connected via a tube with four barriers and two RFID antennas.

### 2.2 Electronics

The MoPSS System consists of an Arduino MKR WiFi 1010 microcontroller with an attached Arduino MKR SD PROTO SHIELD holding a microSD card (Samsung, South Korea) for data collection and control of the RFID reader modules. A small Lithium-Polymer battery is attached to the Arduino with a 3D-printed mount (Supplement file: MoPSS Battery Holder.stl) including a dedicated switch integrated in the housing, to allow disconnecting the battery. Two RFID reader modules (RFIDRW-E-TTL, Priority 1 Design, Australia) and two external antennas (RFIDCOIL-49A, Priority 1 Design, Australia) are used for reading the RFID signals. In order to protect the antenna coils, a support that fitted exactly around the Plexiglas tubes was used, first premade and later self-built using a 3D printer (files available in the Supplements).

The mainboard for the MoPSS system is built on a perfboard and provides the connections between the Arduino and the RFID modules. Three LEDs for visual feedback, three push buttons for user input and reset are added. The mainboard also provides pin header connections for the push buttons, antenna barrel connectors and the power connector (Fig. 2).

**Fig. 2:**
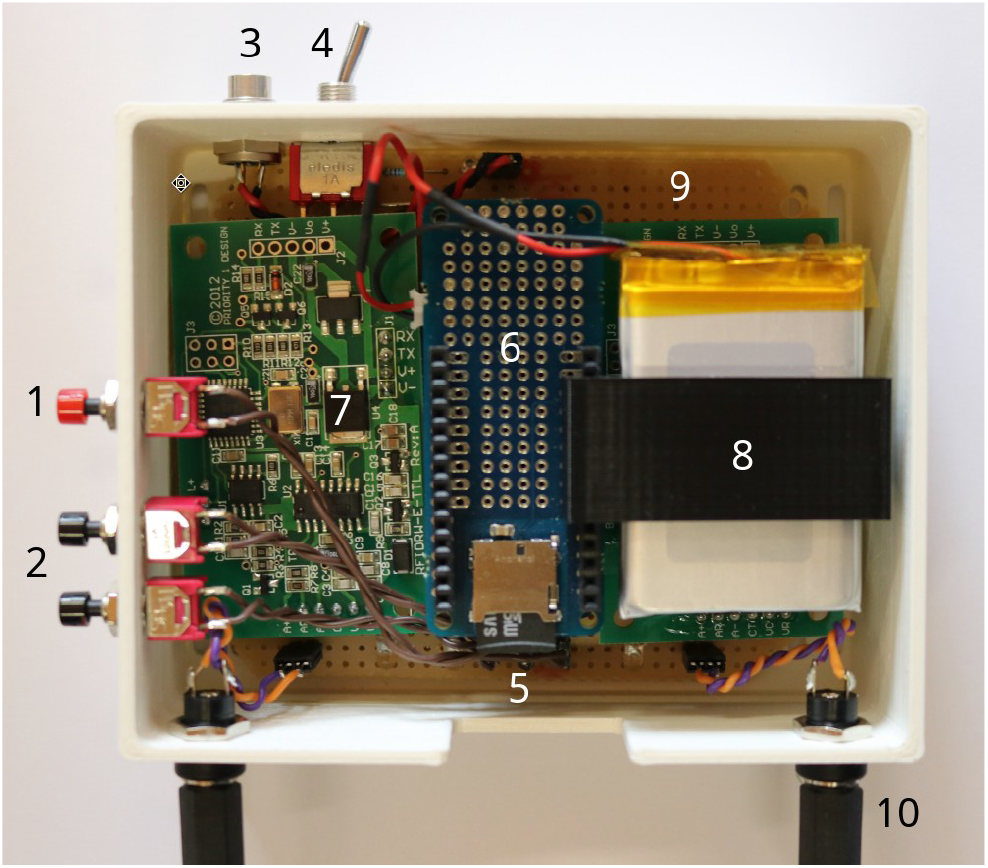
Inner workings of the MoPSS: 1 reset button. 2 button B1 & B2 for user input. 3 power connector. 4 battery on/off switch. 5 microSD card. 6 MKR SD SHIELD, Arduino below. 7 RFID reader module. 8 Lithium Polymer battery with holder. 9 mainboard. 10 antenna connector.

The box for the MoPSS system is printed using polylactic acid (PLA) and consists of a bottom unit with cutout for easy access of the microSD card and mounting holes for the buttons etc. A lid with venting holes for the box is also included (Supplement file: MoPSS Case.stl and MoPSS Lid.stl).

### 2.3 Barrier Construction

Barriers were implemented to slow the mice down while moving through the 31 cm long tube (diameter: 4 cm). To achieve this, we applied four barriers: For both RFID antennas a barrier from below (5 cm from the end of the tube) and a barrier from above (10 cm from the end of the tube) are inserted (see Fig. 3). To install the barriers, 5 mm wide slits have to be cut into the tube. Barriers block about 40 % of the tubes’ diameter and are 4 mm wide. The barriers are made with a 3D printer (Ultimaker 3 Extended, Ultimaker B.V., The Netherlands) using Ultimaker black PLA as material. They are designed with two hooks on either side, so they can be easily inserted into the tube and fixed with a rubber band. The barrier template for the 3D printer is offered (Supplement file: Barrier.stl). In addition, to facilitate the cutting of the tube, a 3D template is provided (Supplement file: Gauge Tunnel Barriers.stl), which assists in drawing exact cutting lines onto the tube.

**Fig. 3:**
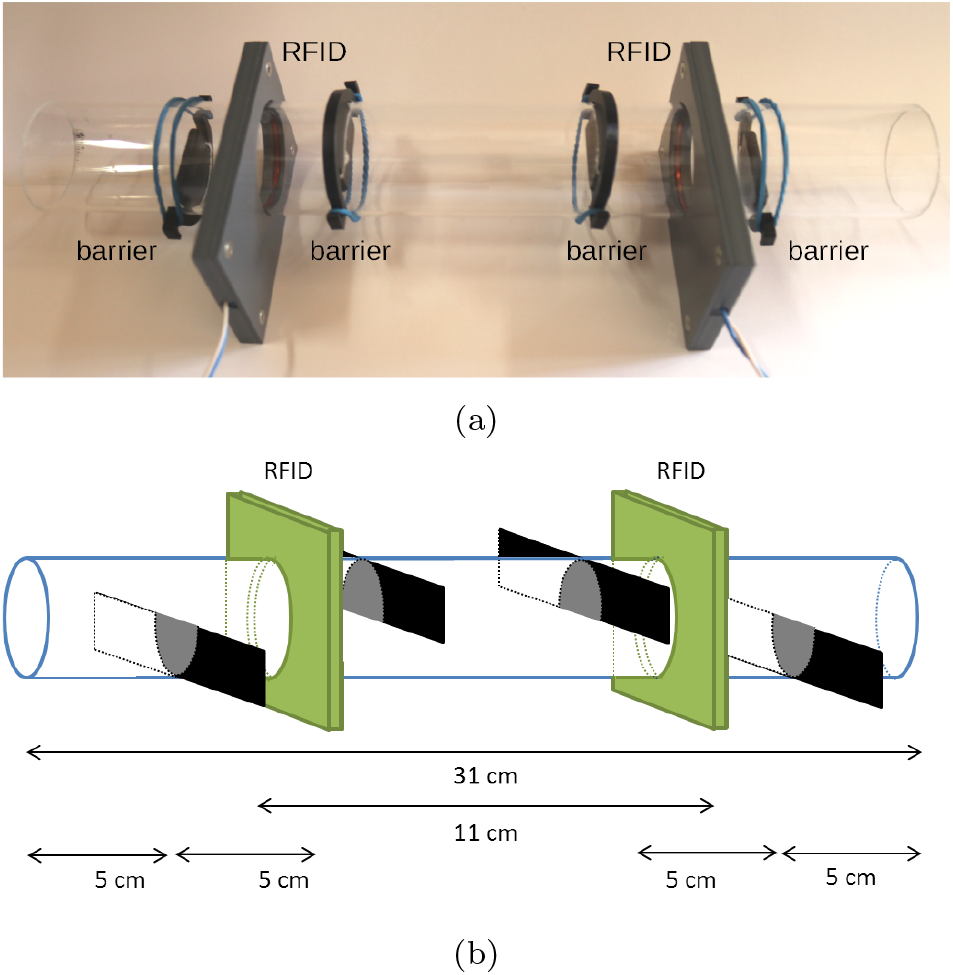
Picture of barrier construction (a) and schematic drawing (b) of barrier construction. RFID: RFID antennas, black: barriers.

### 2.4 Transponders

We use transponders according to ISO 11784/85 (FDX-B transponders, Euro I.D., Germany). The transponder needs to be implanted rostrocaudal for optimal detection sensitivity. The best read performance is achieved when the RFID transponder is oriented lengthwise (0°/180°) to the antenna where read ranges of approximately 4 cm can be achieved. If a transponder were oriented transversely (90°/270°) to the antenna, the read range would approach 0 cm. For more details on the transponder implantation procedure see section *Experiment 1, Animals*.

### 2.5 Software

The Arduino and RFID reader modules run different software each. The RFID modules use proprietary software while the software for the Arduino is available in the appendix. RFID modules: The RFID modules are connected to an antenna each in order to read the unique number of the RFID tag that is within read range and transmit this tag number to the Arduino.

As soon as an RFID tag enters the read range of the antenna, the tag number is read by the RFID module and transmitted to the Arduino. However, the tag number is only transmitted when the tag newly enters the read range.

In order to eliminate interference between the two RFID antennas in close proximity, we decided to enable only one RFID reader at a time for 100 ms, alternately switching between both. As a consequence, every time an RFID reader is re-enabled, any tag it reads will be automatically transmitted because the tag appears as “new” to the RFID reader. This enables us to easily detect when an RFID tag is no longer within the read range of the reader.

Arduino: The Arduino is handling the processing of the RFID tag numbers that are communicated by the RFID modules and adds additional functionality such as visual feedback and logging. Additionally, the Arduino controls charging of the battery that allows coping with short term power loss.

During startup the Arduino connects via WiFi to the internet in order to update the internal Real Time Clock, which is then used during logging to provide accurate timestamps for all RFID tag detections. For the timestamps the unix time is used which is easily processed in further analysis and indifferent to timezones. After successful synchronization the WiFi on the Arduino is no longer required and turned off, thereby greatly reducing power consumption. The battery allows independent operation of the Arduino, guarding the system in case of external power loss for roughly 26 hours. Even though RFID capability is lost while running on battery power, the reader modules will restart without adverse consequences once power is restored. Battery power can also be used for the startup of the MoPSS system at a different location for example if there is no WiFi available inside the animal facility.

The Arduino also controls the LEDs on the mainboard communicating the different states between power on and ready for operation. At the time of writing these are: “searching for WiFi network”, “fetching time from network time protocol server”, “ready for operation”, and “error during setup” indicating a faulty/missing microSD card, inability to connect to the network/synchronize the time. During operation two red LEDs corresponding to the two RFID reader modules are also used to indicate the detection of a tag.

In the event of a successful RFID tag detection, the Arduino saves the data to the microSD card: the antenna number by which the tag was read (A1/A2), the current time (e.g., 1567081062), the tag number (e.g., 900 200000123456) and a flag (E) indicating that this detection corresponds to a mouse entering the read range. When the transponder is no longer detectable, an additional entry is made containing the antenna number, current time, the tag number and the flag X to indicate an exit from the read range. See Table 2 for an example.

**Table 2:** Example of the recorded data provided by the MoPSS.

| Antenna No. | Unixtime | Tag number | Entry/EXit flag |
| --- | --- | --- | --- |
| A1 | 1567081062 | 900_200000123456 | E |
| A1 | 1567081063 | 900_200000123456 | X |
| A2 | 1567081071 | 900_200000123456 | E |
| A2 | 1567081072 | 900_200000123456 | X |

### 2.6 Data Evaluation

Although accuracy of the RFID detections was very high (see section *Experiment 1 Validation, Results*), there were still a few missed detections. We therefore conducted an in depth analysis of the possible combinations of missed detections with the known detections to identify cage changes despite missing data. The resulting R-script can systematically analyse raw data and reliably reconstruct cage changes in the few cases of missing detections. The complete description of this procedure can be found in the Supplementary Material.

## 3 Experiment 1: Validation

In order to compare the accuracy of the MoPSS to manual video analysis we performed a validation experiment using both methods in parallel.

### 3.1 General Procedure

A group of twelve young mice was habituated for 6 days to the MoPSS, including the barrier system in the connection tube before a 24 h video recording was performed. The video recording was then analysed with regard to cage changes and these were compared to the cage changes detected by the MoPSS.

### 3.2 Animals

We chose C57BL/6JCrL mice because it is the most commonly used mouse strain. Twelve female C57BL/6JCrL mice, kept as one group, were used for this experiment. They were purchased in June 2019 at the age of 4 weeks from a commercial breeder (Charles River, Sulzfeld, Germany) and had different mothers and had different nurses to prevent any breeding related effects. At five weeks of age, transponders (FDX-B transponder according to ISO 11784/85, Euro I.D., Germany) were implanted subcutaneously in the neck region (see Fig. 4). In order to prevent potential harm inflicted by the implantation procedure, the mice obtained an analgesic (Meloxicam) the evening before implantation. The transponder injection itself was performed under anaesthesia (Isoflurane) and the RFID transponder was injected directly behind the ears subcutaneously in the neck, so that it was oriented rostrocaudal. After transponder injection mice were placed in a separate cage with bedding and paper for monitoring until they were fully awake again. Then they were returned to their home cage.

**Fig. 4:**
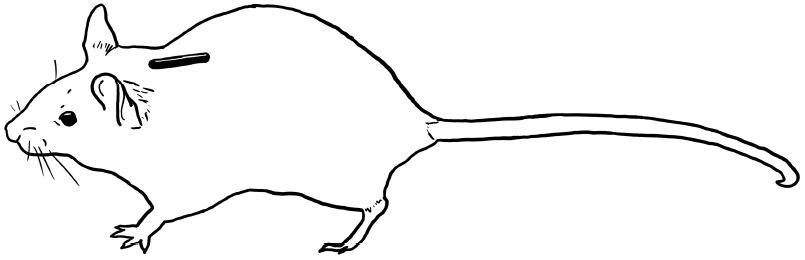
Schematic drawing transponder position. ©Anne Habedank

### 3.3 Housing

In the first weeks, mice were kept in a type IV makrolon cage (L × W × H: 598 × 380 × 200 mm, Tecniplast, Italy) with a filtertop. Food (autoclaved pellet diet, LAS QCDiet, Rod 16, Lasvendi, Germany) and tap water (two bottles) were available ad libitum. The Cage was equipped with bedding material (Lignocel FS14, spruce/fir, 2.5-4 mm, JRS, J. Rettenmaier & Söhne GmbH + Co KG, Germany) of 3-4 cm height, two red houses (The MouseHouse, Tecniplast), four papers, four cotton rolls, 12 strands of additional paper nesting material, and four wooden bars to chew on. The cage also contained a tube (40 mm in diameter, 17 cm long), which was used for tube handling (Hurst 2010, Gouveia 2013).

For the validation of the MoPSS, when the mice were 6 weeks of age, mice were moved into two type III makrolon cages (L × W × H: 425 × 276 × 153 mm, Tecniplast, Italy) with filtertops connected via a perspex tube (40 mm in diameter, 30 cm long) containing barriers from above and below (blocking 40% of the tube diameter with a thickness of 4 mm, see description of barriers in the section *Barrier Construction*). The equipment described above for the type IV cage was equally split unto the two type III cages, except that only one cage contained the handling tube.

Room temperature was maintained at 22 3 ° C, the humidity at 55 ± 15 %. Animals were kept at 12h/12h dark/light cycle with the light phase starting at 8:00 a.m (summer time). Between 7:30 and 8:00 a.m. a sunrise was simulated using a Wake-up light (HF3510, Philips, Germany). Once per week, the home cage was cleaned and all mice were scored and weighed. In this context, mice also received a colour code on their tails (using edding 750 paint markers) to facilitate individual recognition during video recording.

### 3.4 Procedure

With 6 weeks of age, the twelve female C57BL/6J mice were transferred into the test system, consisting of two cages connected with a tube containing four barriers and two RFID antennas (for details see section *Housing* and *Barrier Construction*). After 6 days of habituation to this setup, video recordings of the tube were made for 24 h. To ensure continuous recording of mouse movement, we installed a red light source, which was automatically switched on during the dark phase. The video recordings were conducted with a WebCam (Logitech C390e, Switzerland) using the recording software iSpy 64 (version 7.0.3.0), which automatically cut the videos into blocks of 1 h duration. The WebCam was positioned in a way that ensured a clear view of the connecting tube and the MoPSS, which signalled every RFID detection via two separate LEDs.

Afterwards, we collected the recorded data from the MoPSS and compared the detected cage changes with the 24 h video recordings: We fast-forwarded the video recordings till a mouse was visible and, slowing down the video, monitored then whether the MoPSS signalled via a blinking LED that the RFID tag number of the mouse was detected. In some cases more than one mouse passed through the tube and an additional evaluation whether or not all mice were detected was conducted. Therefore the recorded data from the MoPPS were examined to verify that all RFID tag numbers were recorded at the corresponding timestamp. All missing detections were noted.

As described in Data Evaluation, in addition to just using the data as it was saved by the MoPSS, we also developed a method to improve the received data by means of logical reconstruction (searching the recorded data for inconsistencies in the order of cage changes, for details see section *Data Evaluation*). In the process of evaluating the R script for this logical reconstruction, parts of the video recordings were watched again to compare the results of the script against the true events.

### 3.5 Results

During the 24 hours, 7382 detections were recorded, including 2804 cage changes. On average there are more than twice as many detections as cage changes because mice do not always change cages but sometimes also just stick their nose inside the RFID antenna (poke) and then return to the cage they came from. After the manual comparison of the recorded detections with the 24 h video recordings, we found 9 missed detections, meaning an event in which one of two antennas did not detect the mouse (situation B and C from section Data Evaluation). This led to an error rate of 0.122 % of all the cage changes. There was no cage change detected on video for which both antennas did not detect the mouse (situation D from section Data Evaluation) which would have not been possible to reconstruct due to the missing time stamps. After analysing the data by means of logical reconstruction (as described in section Data Evaluation), we were able to infer the 9 missing detections automatically and correct the corresponding cage changes. In this manner, the error rate was reduced to 0 %.

Analysing the detections, we found that dwelling time between the readers was on average 1736 ms ± 8255 ms, with 87.33 % of cage changes taking ≤ 3 s and 94.27 % taking ≤ 5 s.

### 3.6 Discussion

Validating the MoPSS’ detection with the manual video analysis, we confirmed that the MoPSS reaches a very high accuracy. After logical reconstruction, the MoPSS detection matches to 100 % with the results of the manual video analysis. The only divergence arises in the timestamps (when detections are corrected in their timestamps caused by a situation C as described in section *Data Evaluation* and the Supplementary Material). Here, however, we can assume that the passing mouse was missed by the RFID antenna passed second because it moved too fast out of the antenna’s read range, and therefore, we found that the used timestamps (from the antenna passed first) are only differing by a few seconds from the correct time.

Note that the error rates reported above are only results of one group of mice and thus, they might not be representative for other groups, especially when differing in age, strain or sex. Still, we regard the chosen test group as the optimal one for its purpose: The main difficulty, as explained above, was the velocity of the mice, and that is why we used very young and, thus, fast animals. The mice had six days of habituation to adjust to the new barrier setup. However, it is possible that the mice were not at their highest possible speed. In the study by Bono et al. (2006), it is described that maximum continuous speed increased until day 17 of training for female C57BL/6J mice (10-11 weeks old). Hollinski et al. (2018) described an increase in maximum continuous speed up until week 8 of training. Nevertheless, these studies were conducted on running wheels, whereas for our experiment the maximum speed over a distance of approximately 8 cm in a straight line is the most relevant, as this is the range of the RFID antenna.

We believe our manual video analysis can be considered nearly flawless because, when in doubt, videos were played backwards or in slow motion. This also emphasises the improvement the MoPSS is going to make as an accurate analysis by video was very time consuming.

Comparing the MoPSS’ accuracy to the other available methods for home cage based preference tests (which were described in the introduction) proves difficult. First, accuracy can only be compared to manual analysis, which would make video recordings automatically the most accurate method. However, as we experienced during the development of MoPSS prototypes, especially when using group housed mice, even manual analysis can be complicated. When mice climbed over each other, they were sometimes not distinguishable without the information provided by the RFID antennas.

Second, comparing the MoPSS’ accuracy to other automated tracking systems is in some cases not possible because the studies do not provide any information on accuracy (Linnenbrink and von Merten, 2017; Krohn and Hansen, 2010) or any details on the tracking system except that they used one (Kawakami et al., 2012).

Third, of the remaining two automated tracking systems, the one described by Blom et al. (1992) only uses individually housed mice, which makes data acquisition far easier, but with the disadvantage that the transferability of gained results for group housed mice remains questionable. In addition, Blom et al. (1992) and Tsai et al. (2012) use a correlation between relative dwelling times per cage based either on visual observations or automatically registered cage changes. This, however, does not provide general information on the error rate of the system, it merely states that there is no significant difference between the results. This, however, would change if a cage change was missed after the mice had stayed in this cage for several hours. The paper by Tsai et al. (2012) offers an error rate with 0.26 % of misreported cage changes. In comparison, the MoPSS has an initial error rate (before logical reconstruction which corresponds to RFID reader accuracy) of 0.122 % for missed detections. As explained in the section Data Evaluation, missed detections do not have to lead to a missed cage change when the first RFID antenna the animal was passing through was the one with the missed detection. As only the second RFID antenna reports an actual change in position.

Fourth, it has to be noted that currently no automated tracking system can reach 100 % accuracy at all times (without additional analysis of the data afterwards) because at this time there are situations which can not be identified by automated systems. For example, when a mouse passes through an antenna and another mouse passes the antenna at the same time, two RFID transponders are within the detection range and one RFID tag may obscure the other. However, this is a very rare scenario. Overall, we demonstrated that the MoPPS is equally accurate as video observation and much superior with regard to time taken for analysis.

## 4 Experiment 2: Example data

### 4.1 General Procedure

Experiment 2 is an example of a home cage based preference test conducted with the MoPSS. Two types of bedding material were compared, using one group of twelve mice. The preference test was performed in two consecutive rounds of three days each. In-between rounds, the presentation side of the bedding materials was changed, starting the new round with freshly cleaned cages. The MoPSS was active during the whole duration of the experiment; however, only day 2 of both rounds, respectively was used for analysis.

### 4.2 Hypothesis

We conducted a home cage based preference test, comparing two bedding materials: Pure (cellulose, JRS) and Comfort White (cellulose, JRS). Both bedding materials were known to the mice because they were used before in a conditioned place preference test as the conditioned stimuli. In this test, mice had shown a significant preference for Comfort White bedding during the 10 min habituation as well as during the final test after conditioning. Now, we wanted to investigate whether this preference would persist if mice had not only 10 min but several days of continuous access to the bedding materials.

### 4.3 Animals

Another group of twelve female C57BL/6J CrL mice was used for this experiment. This group was purchased in December 2017 at the age of 3 weeks from Charles River, Sulzfeld. Mice were born by different mothers and had different nurses in order to cope for any possible effects on behavior related to the prenatal and early postnatal phase within the inbred strain. With about five weeks, transponders (FDX-B transponder according to ISO 11784/85, PlanetID, Germany) were implanted under the skin in the neck. The procedure was the same as for the group in experiment 1, except that Meloxicam was given two hours before the procedure instead of the previous evening. In addition, for two mice the transponder implantation had to be repeated at the age of 8 weeks because they lost their transponder immediately after the first implantation.

This group of mice took part in multiple testing of prototypes to develop an automated tracking system. By the time the home cage based preference test was performed to gain example data with the MoPSS, they were around 19 months old. In-between, mice had also participated in other experiments, e.g., T-maze preference tests and conditioned place preference tests (the latter were pre-registered at the Animal Study Registry: Lewejohann, 2024b; Lewejohann, 2024a, the former took place before the launch of the Animal Study Registry).

### 4.4 Housing

Outside experiments, mice were kept in two type IV makrolon cages (L × W × H: 425 × 276 × 153 mm, Tecniplast, Italy) with filtertops connected with a perspex tube (40 mm in diameter), which was equipped in the same way as the two type III cages described for the group in experiment 1.

### 4.5 Procedure

Because this group of mice was usually kept in a home cage system with two connected cages, those cages were identically equipped as always, except that we changed the normal bedding material for different ones: One cage was filled with Pure bedding (cellulose, Arbocel pure, JRS, J. Rettenmaier & Söhne GmbH + Co KG, Germany) and one with Comfort White bedding (cellulose, Arbocel comfort white, JRS, J. Rettenmaier & Söhne GmbH + Co KG, Germany) up to the same height of 3 cm. Both beddings consisted of cellulose, while the usual bedding consisted of conifer wood (spruce / fir). For a picture of the different bedding materials see Fig. 5. The connecting tube was similarly designed as described in Experiment 1, however, we only added barriers from below to facilitate their passing through the tube. This group was older and one mouse was unusually hesitant towards new objects, which had already been observed during several other experiments, and we did not want to exclude it.

**Fig. 5:**
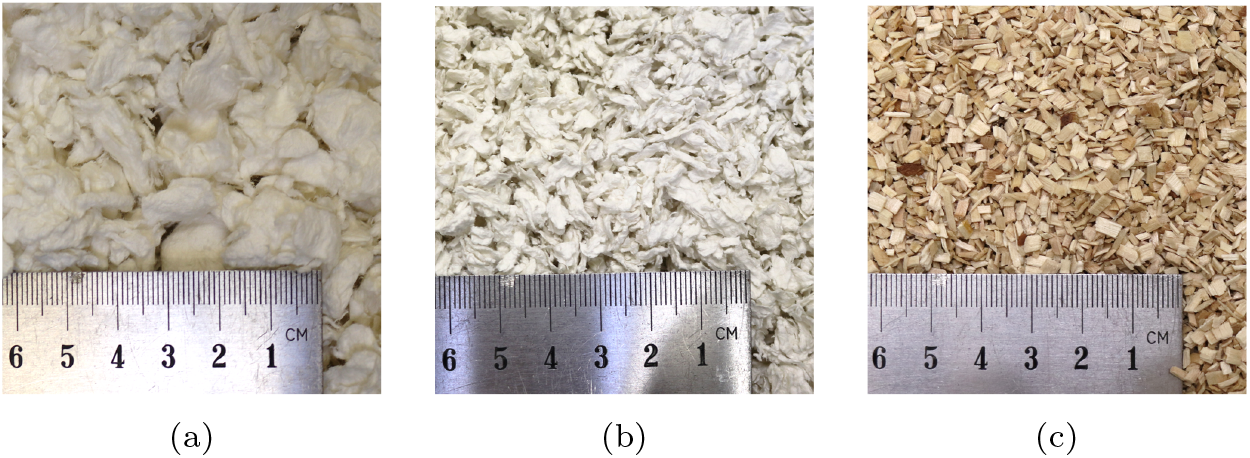
Bedding materials used during the experiment. Comfort White (a) and Pure (b) bedding material were compared in the home cage based preference test and consist of cellulose. (c) The FS14 bedding material consists of spruce/fir chips. This bedding material was not used in the home cage based preference test but was used during normal husbandry conditions.

As it is possible that the spatial position in the room (and its light, noise, room air conditions) influences the preference of the mice (Blom et al., 1992), we performed two rounds, in-between which the presentation sides of the bedding materials were changed. This ensures a discrimination between side and bedding preference. The experiment lasted seven days, with three days presenting bedding material Pure left and Comfort White right (round 1), then switching sides and presenting Pure right and Comfort White left (round 2). On the first day of each round, mice were placed into freshly cleaned and newly equipped cages, placing individual mice alternately into the left and right cage, dependent on the order they entered the handling tube. The first day was considered as habituation day to get the mice accustomed to the new bedding material. The second day was then used for actual data recording. The third day was added for organisational reasons: After approximately 23 h of the third day, the mice were then taken out of the test setup and placed into a separate cage (which contained the spruce/fir bedding they usually had), while preparing the new setup. Mice were then placed into a freshly cleaned and newly equipped cage, this time with changed presentation sides of the bedding. Only the food was maintained; pellets of both cages were mixed and split for the two new cages. The tube connecting the cages as well as the barriers were not cleaned in between. In the second round (just as in the first round), only the second day was analysed, leaving the first for habituation.

### 4.6 Statistical Analysis

During the preference tests, RFID detections by the two RFID antennas were automatically saved by the Arduino onto a microSD card. Each record included a timestamp (synchronized before the start of the experiment via internet connection), antenna number (A1 or A2) and the detected RFID tag number. With the help of R studio (Version 1.1.383, requiring on R 3.0.1+), the data recorded by the Arduino was analysed for missing detections (see section *Data Evaluation*). Following this procedure cage changes were extracted. In the case of missing detections, in which one RFID antenna did not detect the cage change, the timestamp of the detection of the second antenna was used, arguing that the missing detection resulted from a mouse passing too fast through the tube, which should lead to a roughly similar detection timestamp for both antennas. We decided against subtracting the time spent in the tube from the stay duration. Thus, we calculated stay times for each mouse in each cage as times between cage changes when a mouse entered a new cage (only detections by the antenna passed second).

For each mouse, stay times in each cage were then summed up per day. As already mentioned, we analysed only the second day of each round because the first day was considered habituation time. Thus, for the investigated 48 h, the percentage of time spent in each cage was calculated for each of the twelve mice. These percentages were then used for further analysis to compare side preference (left vs. right cage) and bedding preference (Pure vs. Comfort White, whereby presentation sides were switched after the first round). To test for normal distribution, the Shapiro-Wilk test was performed in R. The data was considered normally distributed (p > 0.05); therefore, a t-test was used to compare the stay time percentages with a chance level of 0.5 (the expected relative stay time if mice had no preference for one of the two cages). In all statistical tests, significance level was set to 0.05, and result values are given as mean and standard deviation.

### 4.7 Results

During the two analysed days, mice changed cages between 52 and 178 times per 24 h (100.75 ± 31.84 cage changes). Comparing the times the twelve mice spent in the two cages, we found that during the whole experiment mice stayed significantly longer in the right compartment, namely 57.49 +/3.83 % of the time (t(11)= −6.77, p < 0.001, see Fig. 6a). For the different bedding materials, on the other hand, there was an even clearer preference: Mice stayed 72.76 ± 3.00 % of the time in the compartment with Comfort White bedding (t(11) = −20.19, p < 0.001, see also Fig. 6c).

**Fig. 6:**
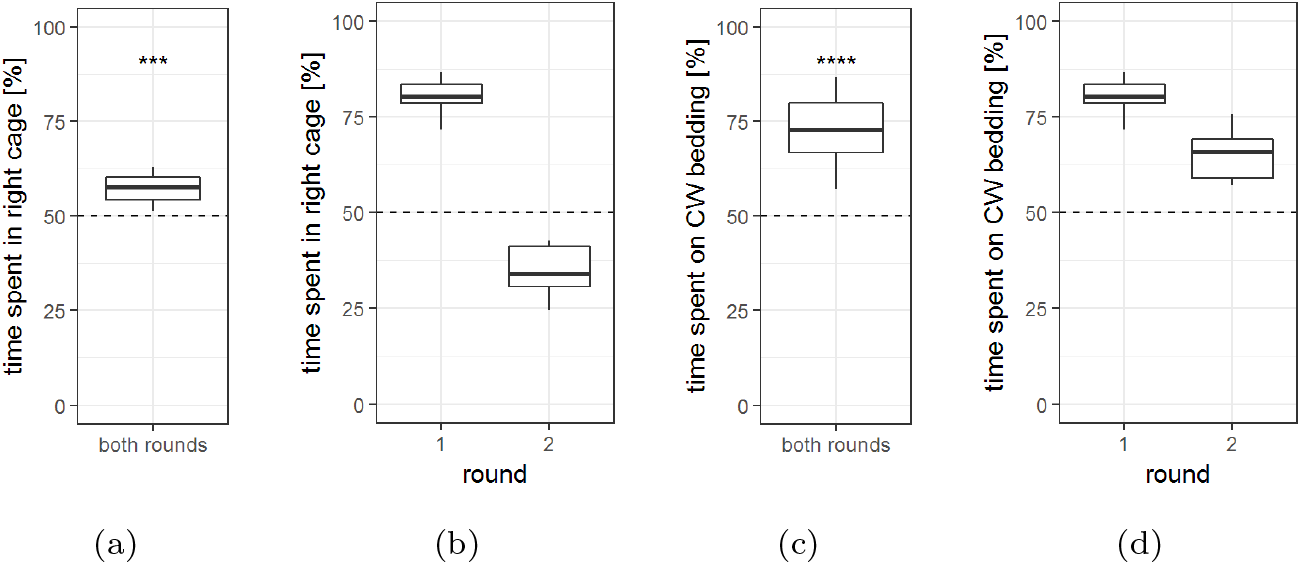
Time spent (%) in the two cages, analysed by cage side and bedding material. Time spent in the right cage (a) in total (48 h), or (b) with regard to round (24 h). Time spent in the cage with the Comfort White bedding material (c) in total (48 h), or (d) with regard to round (24 h). Comfort White bedding material was presented in the right cage during the first round and in the left cage during the second round. CW = Comfort White *** p < 1×10^−4^, **** p < 1×10^−9^ t-test comparison to chance level, n = 12

### 4.8 Discussion

In this experiment, stay times of twelve mice on Comfort White and Pure bedding material were compared, whereby stay time was only analysed after one day of habituation, and the presentation side of the bedding was changed in-between to control for side preference. When looking at Fig. 6b, which compares the side preference on the second day of both rounds, side preference seems to be more distinct during the first round than the second. This was also reflected in a significant side preference, which could be due to spatial reasons (position in the room etc., Blom et al., 1992). Another explanation could be that the condition preference (for the bedding material) changed over time, becoming less strong and thus, leading to a side preference when compared with the round before.

Nevertheless, mice had a distinct preference for the cage with Comfort White bedding compared to the cage with Pure bedding. Thus, during this home cage based preference test we could confirm the results already obtained during the two ten minute observations of the conditioned place preference test: Comfort White bedding is preferred over Pure bedding by this group of twelve C57BL/6J mice.

The main purpose of this experiment was to test the new setup in a weeklong experiment as well as to validate the bedding preference previously observed during a conditioned place preference (CPP) test. We have to emphasize that the result of this preference test can not be generalized for C56BL/6J mice: Although we tested the preference of twelve mice, they were all together as one group in the test system and, thus, might be considered as only one independent sample. Indeed, it is possible that the mice influenced each other in their stay a) by the behaviour of dominant mice, b) by avoiding or following individual mice, c) by preferring to not sleep alone over individual bedding preferences. As stated above, the bedding material was also familiar to the mice and as it was presented first in an experimental environment, it is possible that this might have had an influence. Thus, this test would have to be repeated with more groups and also younger mice for a more generalized conclusion. In any case, the preference test was successful in showing the feasibility of the MoPSS under the experimental conditions of a home cage based choice test.

## 5 Conclusion

In this paper, we offer the construction description to build an automated tracking system, which can be used to facilitate the analysis of home cage based preference test. We showed that the MoPSS is accurate even for fast mice and its error rate can be further reduced close to 0 % with the help of additional logical reconstruction of the data. We also presented an example experiment with the corresponding results, in which we compared two different bedding materials.

With this automated tracking system, analysis of home cage based preference tests will become much easier: They will be less expensive, require less time for the data analysis, and will have much finer data resolution. The MoPSS is able to track individual mice and, therefore, it is suitable for group experiments. In our laboratory the MoPSS is already being used to compare multiple enrichment conditions with regard to the mice’s preference over several months. We want to emphasize the great advantages of the MoPSS to existing systems: It is even able to detect fast animals and can be easily rebuilt. Currently, we are working on a further improved version with an RFID reader module without proprietary software and increased detection rates. In addition, in the near future we will be adapting the MoPSS system to be suitable for larger animals such as rats and guinea pigs that require a tube diameter of more than 4 cm. On the basis of the construction description, it is also possible to adjust the MoPSS to other research questions. For example, we are working with a modified MoPSS onto which automated doors and levers or nose poke sensors can be added to test not only for preference but also for the strength of preference by letting the animals work for the access to the other cage (Lewejohann and Sachser, 2000; Sherwin and Nicol, 1996; Sherwin and Nicol, 1995). Using only one RFID antenna, the MoPSS can also be used to record activity data in the home cage. In addition, the MoPSS might also be used to study group dynamics and the influence of individual group members on the position of the whole group.

## Supplementary Material

Supplementary Material such as 3D printing templates, Arduino code, the R evaluation script and raw data of the validation experiment can be found under: https://seafile.bfr.berlin/d/5045377fc7694df5b7a4. Passwort: mousemouse If there is interest in the video recordings of the validation experiment, please contact us via email.

## Ethical approval

All experiments were approved by the Berlin state authority, Landesamt für Gesundheit und Soziales, under license No. G 0182/17 and were in accordance with the German Animal Protection Law (TierSchG, TierSchVersV).

## Acknowledgements

The authors thank the animal caretakers, especially Carola Schwarck, for their support in the animal husbandry.

