## Supplements for "O mouse, where art thou? The Mouse Position Surveillance System (MoPSS) - an RFID based tracking system"

#### 1 Glossary

Before going into details, we will have a short definition of the wording used here:

**detection:** When a mouse moves through the tube, the RFID tag number of its transponder is detected by the RFID antenna. The RFID reader connected to the RFID antenna transmits the tag number to the Arduino, which can then save it onto the microSD card.

**RFID antenna:** To simplify the explanation, we will use "RFID antenna" synonymous to "RFID reader". Note that "first antenna" is always referring to the first antenna the mouse passes through when moving to the other cage, independent from direction. In the following sections, we will also refer to it as "A1", irrespective of its position (left or right). In the same way, "second antenna" (A2) is referring to the second antenna the mouse passes through and consequently the antenna that is closer to the new cage.

**mouse:** Technically, only the RFID tag number of the mouse's transponder is detected by the RFID antenna. However, we will speak of "mouse".

**cage change:** Cage changes are determined by consecutive detections on both antennas, as caused by a mouse changing cages and consequently passing first A1, then A2. A "cage change" is synonymous to "side change", "passage"

Anne Habedank · Birk Urmersbach · Pia Kahnau · Lars Lewejohann
German Federal Institute for Risk Assessment (BfR), German Center for the Protection of Laboratory Animals (Bf3R), Diedersdorfer Weg 1, D-12277 Berlin, Germany


Lars Lewejohann
German Federal Institute for Risk Assessment (BfR), German Center for the Protection of Laboratory Animals (Bf3R), Diedersdorfer Weg 1, D-12277 Berlin, Germany
Institute of Animal Welfare, Animal Behavior and Laboratory Animal Science, Freie Universität Berlin, Königsweg 67, D-14163 Berlin, Germany

or “transition” used in other studies.

### 2 Data Evaluation

During recording, RFID detections were automatically saved onto a microSD card by the Arduino. Each detection includes a timestamp (synchronized before the start of the experiment via an internet time server), antenna number (A1 or A2), and the unique RFID tag number of the mouse. The recorded data is then analysed for each mouse individually by identifying cage changes. However, if a mouse while changing between cages is not detected by one or by both RFID antennas, the simple approach of looking at consecutive detections does not work anymore. In the following, we will explain how to handle these situations by deducing the mouse position from the available data.

#### 2.1 Four situations for data acquisition

Four situations can be distinguished. (A schematic drawing for the following explanations is depicted in Fig. 1.)

A) **Both antennas detect the mouse**, this is the common/regular case.

B) **The first antenna (A1) does not detect the mouse, but the second antenna (A2) detects the mouse (~~A1~~ → A2).** In this case, the cage change is easily deductible since the mouse must have passed the first antenna in order to get to the second. The missing information is the point in time when the mouse passed/entered the first antenna.

C) **The first antenna (A1) detects the mouse but the second (A2) does not (A1 → ~~A2~~).** Here, the cage change is not immediately identifiable because a detection of the mouse on the first antenna (A1) does not necessarily indicate a cage change. Indeed, dwelling in the range of the antenna without completely passing through the tube is occurring commonly (see Tab. XX). The fact that the mouse has passed the tube is becoming obvious the next time the mouse returns and passes again through the antennas in reverse direction (A2 → A1). We know from observations, that mice usually do not spend prolonged time within the tubes (98.82 % of cage changes in the validation experiment took $\leq$ 10 s). Therefore, it can be inferred that a cage change must have taken place earlier when the mouse is detected at antenna 2. Now that we know that we have missed a cage change, we can look at the previously recorded data and infer when this cage change most probably has happened. For this we use the timestamp of the last mouse detection at A1. This is the best approximation as to when the cage change happened.

D) **Both antennas do not detect the mouse (~~A1~~ → ~~A2~~).** In this case, there is no inference possible because there is no information on the time when the missed cage change happened, apart from the general time frame between two successful cage changes.

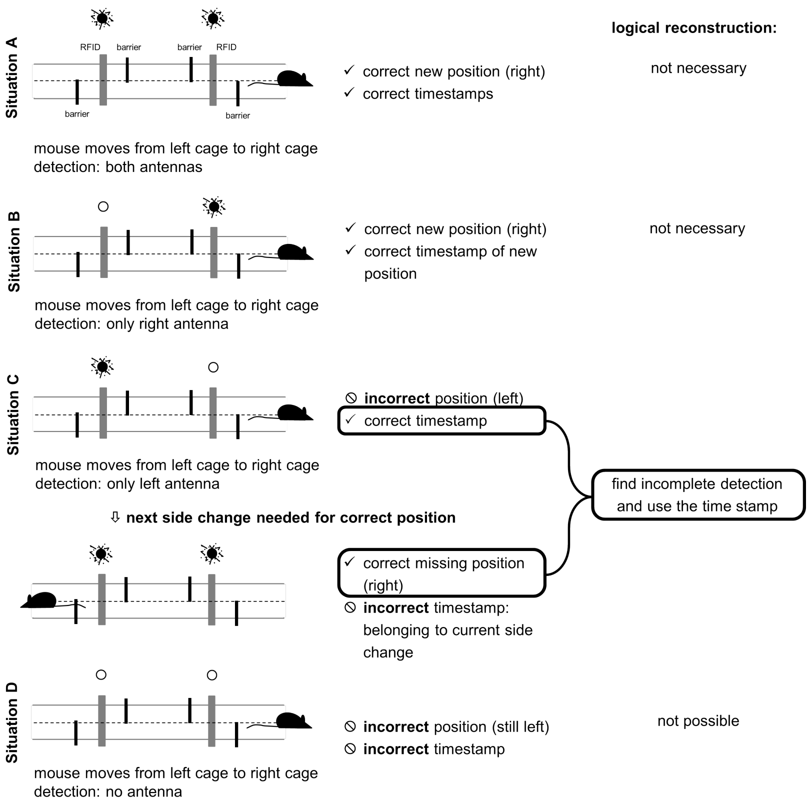


Fig. 1: Four possible situations that might arise during cage changes and how to reconstruct the actual cage changes from them. For a more detailed description of how these situations are handle by the R script, see Fig. 2.

### 2.2 Handling situations A-D

Our main focus was first, to find the cage changes in which one RFID antenna did not detect the mouse (B and C), and to correct possible false timestamps (C, as the time belongs to a new cage change when from the antenna's perspective the mouse appears for the first time on this side), and second, to identify cage changes which were completely missed by the antennas (D, wherever possible, as explained above). To achieve this, we developed

an R script to help with the logical reconstruction of the data (available here: https://seafile.bfr.berlin/d/5045377fc7694df5b7a4, Passwort: mousemouse). At the end, in Table 1 it is shown how the output of the evaluation script of Experiment 1 (validation) looks like.
Our dataset contains a timestamp and the antenna number where the detection occurred. Apart from cage changes, around 62 % (see [Data1]) of our data points consist of detections we considered "pokes". These are detections in which a mouse is recorded (multiple times) at the antenna without passing through it. This is due to dwelling near the beginning of the tube.
In short, the procedure of the R script is as follows (also depicted as a schematic drawing in Fig. 2):

1) Whenever a mouse was detected by an antenna by which it was not detected before, this was identified as a cage change and labelled according to its duration: time passed between detection by the first antenna and the detection by the second antenna. Based on observations made in previous tests, cage changes including detections at both antennas within 3 s were assumed as safe.

2) On the basis of the dataset with safe cage changes, we now looked for two consecutive safe cage changes and subsequently examined all detections in between these two cage changes.

2a) If the two safe cage changes were impossible, e.g., the mouse changed from left to right and again from left to right, the detections in-between were examined. This leads to three possible outcomes:
First, if there was no detection at all between the two safe cage changes, both RFID antennas must have missed the mouse (Fig. 1 D).
Second, if there was only one detection between the two safe cage changes, one of the two antennas must have missed the mouse (Fig. 1 B or C). As we know that an undetected cage change must have happened between the two safe cage changes, we used the timestamp from the single detection and reconstructed the missing cage change. Since cage changes are usually fast, we decided to accept the introduced uncertainty of a few seconds.
Third, if there was more than one detection between the two safe cage changes, we looked for cage changes lasting longer than 3 s. If there was only one such cage change, it was regarded as true, most likely resulting from a B or C situation depicted in Fig 1.

2b) If the two safe cage changes were possible, we examined the detections in-between for occurrence of detections indicating additional cage changes (i.e., presence detection in the wrong cage).
These could be caused by, for example, two cage changes during which one of the antennas did not detect the mouse. E.g., a mouse moves from left to right cage and is detected by second antenna (situation C), moves back from right to left cage and is again detected by second antenna (situation C) (~~A1~~ $\rightarrow$ A2, ~~A2~~ $\rightarrow$ A1). Thus, these two cage changes lead to a detection first left and then right, which would resemble a cage change from left to right. As a result, presence would be assumed in the wrong place, not

matching the detected cage changes before and after. This mismatching is a first criterion, when finding these situations. As an additional criterion, the detections by the two antennas (originally from two cage changes) have to be more than 3 s apart, so that the mouse had time to leave the tube in-between cage changes and before passing again through the antennae.

3) After looking at possible and impossible cage changes, we went through the whole data set of each mouse again, to find additional cage changes which might have not fallen into the previous categories. For example, if between added cage changes were additional detections in the other cage which were not explained by a cage change yet (see Fig. 2b), this would be detected now. To do so, we again examined the detections in-between the now secured cage changes: Were there more than two detections which indicated a cage change (= the mouse was detected by a different antenna then before)? If so, and the time passed between the detections was under 15 s, we assumed that this was also a real cage change, but one in which the mouse moved slower than usual. (This was caused, for example, by multiple mice in the tube, which blocked each other's way.) As a test, we then also included cage changes taking even longer than 15 s, and this also proved to be correct, when comparing them to the video recordings (see following section).

### 2.3 Edge cases of the Evaluation

**Situation D two times in a row:** Although extremely rare, it is possible that a mouse is missed by both antennas when passing through the tube (situation D). In principle this could happen two times in a row leading to an undetectable error based on evaluation of the order of cage changes. E.g., a mouse moves from the left to the right cage, then two cage changes are missed, and the next seen cage change happened logically reasonable from the right to the left cage. However, we could show that situation D (both antennas were missed) is very unlikely and therefore, it is even more unlikely that this occurs two times in a row.

**Situation C followed by situation B:** When a mouse passes from one cage to the other and is not immediately detected by the antenna corresponding to the new cage (A1 $\rightarrow$ ~~A2~~), this error can be inferred from the next regular cage change (situation C). However, this correction is not possible when during the subsequent cage change the same antenna (now A1, formerly A2) does not detect the mouse (~~A1~~ $\rightarrow$ A2). In this case, the recorded data will provide no hint that the mouse has been in the other cage. However, we did not observe this at all during evaluation and thus deem this situation to be very unlikely.

#### 2.4 Customization of the evaluation script

Depending on the research question, the evaluation script can be freely customized. The script is well commented and easy to apply for anyone. See the script and our dataset (in the appendix/online) to try the evaluation first hand.
For setups in which the time to change cages for the animal is shorter or longer than the default of three seconds (e.g., if the distance between antennas is longer), the time for a safe cage change can easily be adjusted. If the absolute highest certainty for cage changes is needed, all detections which were deduced from missed antenna detections can be removed from the dataset (loss of 3.98% cage changes for the validation dataset of Experiment 1). There is no limit to the number of animals or duration of the experiment.

Table 1: Output from evaluation script of the validation experiment (Experiment 1). Detection and cage change are defined as described at the beginning in the glossary. Percentages are calculated as part of the total cage changes each mouse made.

| | Mouse Number | | | | | | | | | | | | | |
|---|---|---|---|---|---|---|---|---|---|---|---|---|---|---|
| | 1 | 2 | 3 | 4 | 5 | 6 | 7 | 8 | 9 | 10 | 11 | 12 | Total | Percentage |
| Error 1 | 0 | 0 | 4 | 1 | 0 | 0 | 1 | 1 | 0 | 0 | 0 | 0 | 7 | 0.095 |
| Error 2 | 24 | 14 | 10 | 11 | 13 | 22 | 13 | 15 | 9 | 12 | 8 | 11 | 162 | 2.195 |
| Error 3 | 0 | 0 | 0 | 0 | 0 | 0 | 0 | 0 | 0 | 0 | 0 | 0 | 0 | 0 |
| Error 4 | 8 | 0 | 8 | 12 | 9 | 13 | 6 | 2 | 7 | 34 | 3 | 23 | 125 | 1.693 |
| Error 5 | 0 | 0 | 0 | 0 | 0 | 0 | 0 | 0 | 0 | 0 | 0 | 0 | 0 | 0 |
| Events | 997 | 582 | 599 | 552 | 500 | 873 | 431 | 722 | 808 | 569 | 327 | 422 | 7382 | 100 |
| Transitions | 398 | 220 | 247 | 211 | 182 | 319 | 148 | 319 | 289 | 205 | 125 | 141 | 2804 | 37.984 |

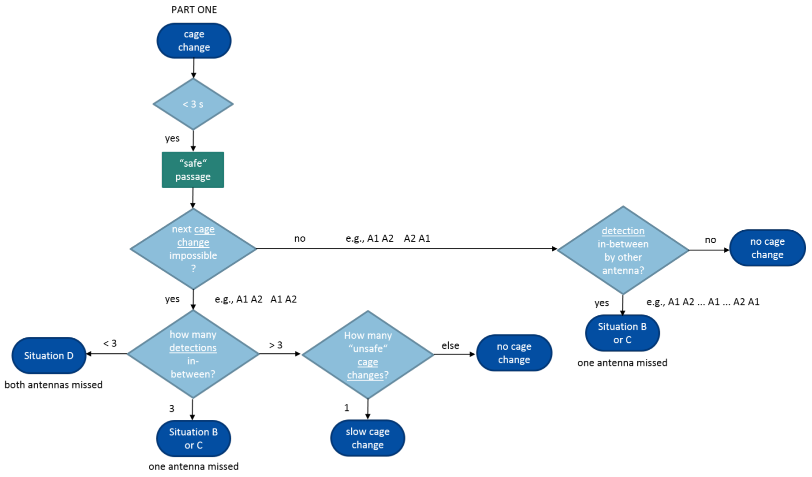


(a)

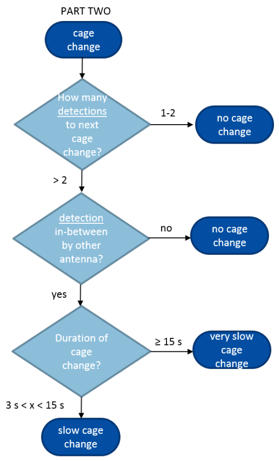


(b)

Fig. 2: Logical reconstruction with the help of the R script. A) Part one: First, cage changes are identified and "safe" cage changes are defined based on their duration (< 3 s between both antenna detections). Next, the cage changes are compared to the following cage change and depending on whether this sequence is possible or impossible, the detections in-between the two cage changes are analysed. B) Part two: The data set with safe cage changes is recalculated, including the reconstructed cage changes from part one. Then, the recorded data are examined for additional cage changes, indicated by a detection from the other antenna in between.
